# m^6^A pathway suppression reprograms gene expression during the integrated stress response

**DOI:** 10.64898/2026.08.05.742933

**Authors:** Shino Murakami, Samie R. Jaffrey

## Abstract

The integrated stress response (ISR) orchestrates cellular adaptations through translational repression and upregulation of ATF4, a transcription factor. However, ATF4 only accounts for a minority of ISR-induced gene-expression changes. Here we show that the ISR reshapes the transcriptome by stabilizing *N*^6^-methyladenosine (m^6^A)-containing mRNAs that are normally unstable. We show that mRNA stabilization, not transcription activation, explains much of the transcriptome response during the ISR, and that this stabilization selectively occurs on m^6^A-mRNAs. m^6^A-mRNA degradation is a translation-dependent process and translation repression during the ISR stabilizes m^6^A-mRNAs. Consequently, the ISR leads to increased m^6^A-mRNA expression, thereby upregulating their protein output despite overall diminished translation. Notably, stabilization of m^6^A-mRNAs by m^6^A depletion is sufficient to induce transcriptomic and proteomic features of amino acid-depleted cells. Together, our study identifies the direct coordination of translation suppression and transcriptome reprogramming through m^6^A during the ISR.

## MAIN

The ISR is a major pathway by which cells sense diverse types of stress and induce gene expression changes that promote cellular homeostasis and survival^1,2,3,4,5,6,7^. A key step in the ISR is stress-induced phosphorylation of eIF2*α*, which impairs translation initiation^8,9,10,1,11,12,13,14^, and increases expression of ATF4, a transcription factor that induces the expression of genes linked to cellular survival responses^2,15^. However, ATF4 only accounts for ∼7.5% to ∼36% of the gene expression changes seen after the ISR induction^16,17^. It remains unclear which additional pathways are activated by the ISR that account for ATF4-independent transcriptomic responses to stress.

A major form of cell stress that is sensed by the ISR is amino acid deprivation. Amino acid depletion leads to activation of GCN2 (general control nonderepressible 2), a kinase that phosphorylates eIF2*α*^11^, leading to reduced translation levels^18^. In amino acid-depleted states, the ISR leads to altered gene expression patterns which allow the cell to conserve resources, enhance nutrient acquisition, and promote stress resistance^19^. Thus, the ability of the ISR to alter gene expression is critical for cellular adaptation to cell stress.

Many of transcripts that are upregulated during the ISR, such as *Mdm2, Atg5, and Atg7*, contain high levels of *N*^6^-methyladenosine (m^6^A), a modified ribonucleotide that is major mediator of mRNA instability. m^6^A is enriched in mRNAs that encode developmental regulators^20,21^ and proteins that contribute to cell stress response pathways^20,22^. These transcripts include metabolic regulators (e.g. *Scd1*, *Dgat2*, *Smpd3, PPARA*),^23,24,25^ DNA-repair pathway proteins (e.g. *Mdm2*, *Rgs12, Polk*)^26^, and autophagy genes (*TFEB*, *RB1CC1*/*FIP200, Atg5, Atg7*)^27,28,29,30,31,32^. Under basal conditions, m^6^A mRNA degradation ensures the low expression of these transcripts. Indeed, many studies have shown that m^6^A is required for proper cellular responses to diverse types of cellular stress^27,33,34,35,36^. These findings raise the possibility that suppression of the m^6^A-mRNA degradation pathway during the ISR may account for the global mRNA stabilization and thus the increased expression of these transcripts.

m^6^A-mediated mRNA degradation depends on active translation^37,36,38^ and can be affected by pathways that regulate translation levels^36^. Notably, a key step in the ISR is phosphorylation of eIF2alpha, which reduces cellular translation rates. Here we show that a large fraction of the gene expression changes induced after induction of the ISR are mediated by global suppression of m^6^A-mediated mRNA degradation. Although the ISR is known to be associated with induction of ATF4-dependent transcription, we show that the ISR also causes global mRNA stabilization, which has a major contribution to the overall transcriptomic alterations seen in the ISR. We find that the mRNAs that are most stabilized are those that are most enriched in m^6^A, and their stabilization is due to the reduced translation seen during the ISR. We find that ISR-mediated stabilization of highly methylated mRNAs accounts for much of the altered transcriptomic and proteomic patterns normally seen during the ISR. Thus, these results show that suppression of m^6^A-mediated mRNA degradation is a key component of the ISR and the cellular adaptation to stress.

## RESULTS

### The ISR leads to increased transcript abundance despite reduced transcription rates

Gene expression changes during the ISR are thought to be mediated by transcriptional activation^39,40^. The transcription factor ATF4 is the master regulator of transcriptional changes during the ISR^39,40^. However, a large part of gene expression changes are still seen in ATF4 knockout cells^16,17^. We therefore wanted to more fully understand the diversity of gene expression changes during the ISR. We used TimeLapse-seq, which involves metabolically labeling cells with 4-thiouridine (4sU) to monitor both RNA stability and transcription rates^41^.

TimeLapse-seq demonstrated clearly increased transcription rates of ATF4-dependent gene expression, including *Ddit3*, *Herpud1*, and *Sqstm1* (**Fig. 1a**), after amino acid deprivation in mouse embryonic fibroblasts (MEFs). Unexpectedly^39,40^, we also found globally suppressed transcription rates. Of the 10,713 transcripts that were measured in TimeLapse-seq, 3,823 genes showed ≥0.5-fold decrease in transcription rate, while only 335 genes exhibited increased transcription by this amount (**Fig. 1b**). There results suggest that the ISR is generally associated with decreased transcription, despite the increase in transcription of ATF4 target genes.

**Fig. 1:**
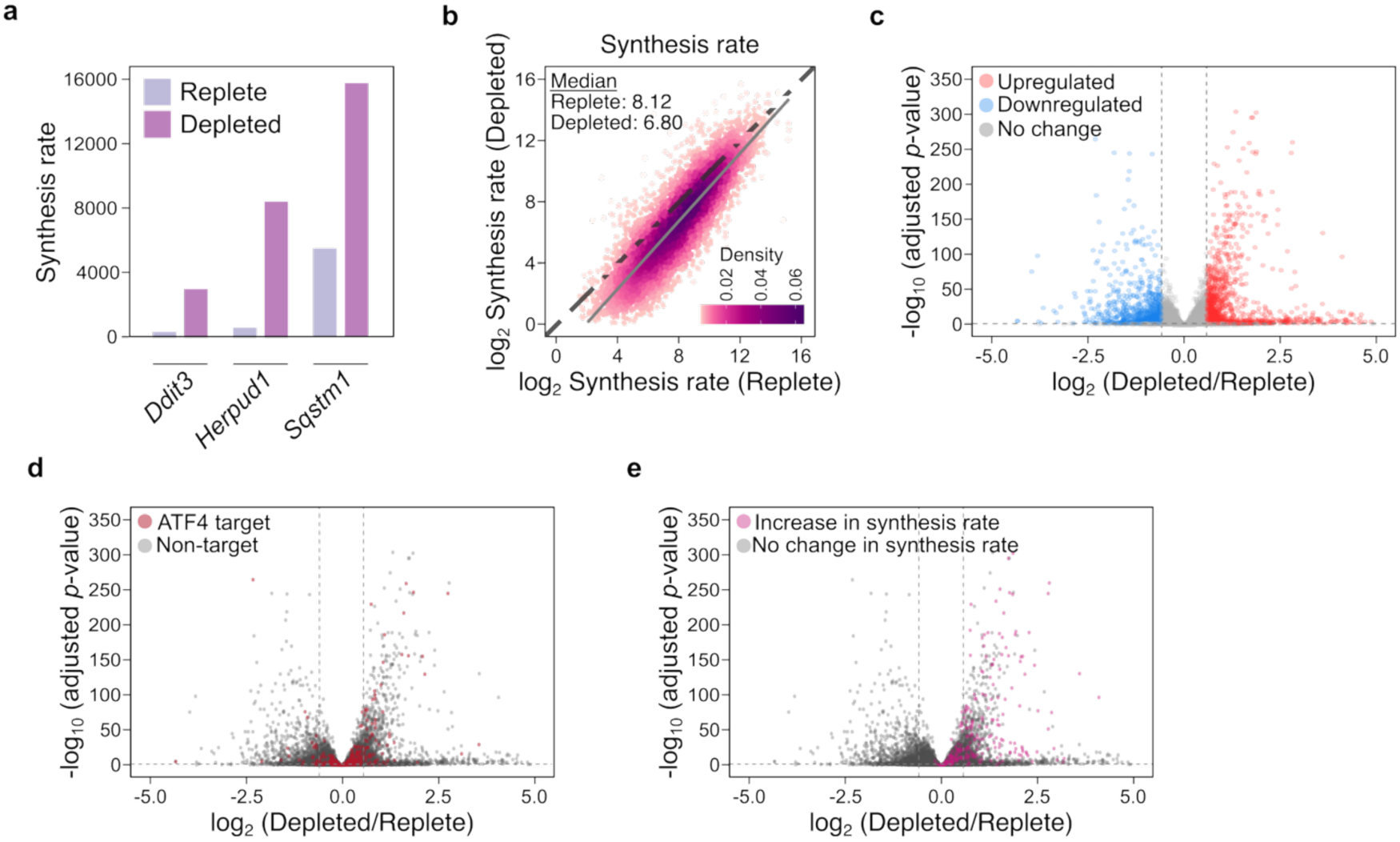
Amino acid depletion globally reduces mRNA synthesis. (a) Amino acid depletion induces increased mRNA synthesis of known ATF4 target genes *Ddit3* (Chop), *Herpud1, Sqstm1* (*p62*)^79^. Shown are the mRNA synthesis rates in amino acid-replete and amino acid-depleted states for known ATF4 target genes. Synthesis rate is defined as an arbitrary unit of RNA produced for a given gene within a given time^73^. (b) Amino acid depletion globally reduces mRNA synthesis rates. mRNA synthesis rates were plotted for transcripts obtained from amino acid-replete and amino acid-depleted wild-type MEFs. Synthesis rate is defined as an arbitrary unit of RNA produced for a given gene in a given unit of time^73^. mRNA synthesis rates were generally reduced following amino acid depletion. *n* = 10713 transcripts. *n* = 3 biological replicates for each condition. (c) Amino acid depletion increases expression of large subset of genes. Changes in steady-state mRNA expression after amino acid depletion in MEF were plotted. Red and blue indicate genes upregulated and downregulated (>0.5-fold change, adjusted *p*-value < 0.05). *n* = 10713 transcripts. *n* = 3 biological replicates for each condition. (d) Same plot as in panel (c). In this plot, ATF4 target genes defined by ATF4 ChIP-seq^75^ are shown in red and all other genes are plotted in gray. (e) Same plot as in panel (c). In this plot, genes that are transcriptionally activated during the ISR are shown in pink and all other genes are plotted in gray.

We also examined gene expression by RNA-seq, expected a similar overall trend as seen in TimeLapse-seq. For our RNA-seq experiments, we used differential gene expression, comparing amino acid replete and amino acid-depleted MEFs. We asked if the RNAs that were transcriptionally suppressed in TimeLapse-seq are downregulated for their relative abundance in RNA-seq. However, we did not find this effect. Instead, we found an overall similar number of genes that were downregulated and upregulated in their relative abundance in RNA-seq (912 genes and 1228 genes, respectively, >0.5-fold change, adjusted p-value < 0.05) (**Fig. 1c**, **Supplementary Table 1**). As expected, some of the upregulated transcripts were known ATF4-target genes. However, most of the upregulated transcripts were not ATF4 targets (**Fig. 1d**). Furthermore, many of the upregulated transcripts in the steady-state levels measured by RNA-seq surprisingly showed no significant increase in transcriptional rates in the TimeLapse-seq dataset (**Fig. 1e**).

These results suggest that many transcripts show relatively increased mRNA abundance despite generally reduced transcription rates.

### The ISR induces widespread mRNA stabilization

Since transcription is broadly suppressed after amino acid depletion, we asked whether the increase in mRNA abundance could be due mRNA stabilization. To test this, we used the TimeLapse-seq dataset to measure mRNA half lives. We found a widespread increase in mRNA stability after amino acid depletion (**Fig. 2a**). Of the 10,713 transcripts detected, 4,349 transcripts showed >0.5-fold increase in mRNA half-life, with some achieving >3-fold increase in half-life. Only 6 transcripts were destabilized by >0.5-fold (**Fig. 2a**). These results suggest that global mRNA stabilization could account for the increased transcript levels during the ISR (**Fig. 2b**).

**Fig. 2:**
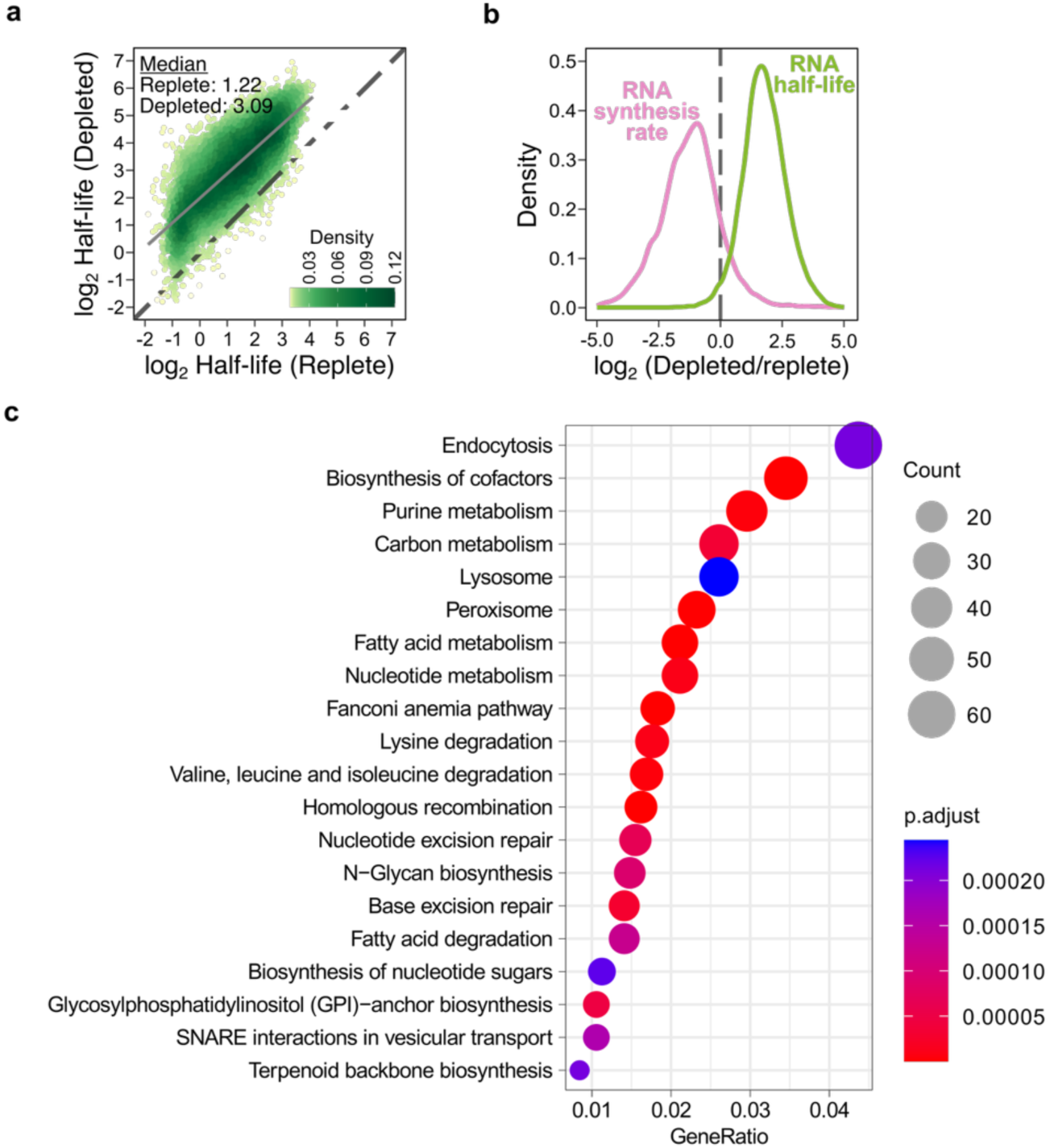
mRNA stabilization is a major feature of the transcriptome-wide response to amino acid depletion. (a) Amino acid depletion globally increases mRNA half-life. mRNA half-lives were plotted for transcripts obtained from amino acid replete and amino acid-depleted wild-type MEFs. mRNA half-life was globally increased after amino acid-depletion. *n* = 10713 transcripts. *n* = 3 biological replicates for each condition. (b) Amino acid depletion globally reduces mRNA synthesis rates and increases mRNA half-life. mRNA synthesis rates and half-life under amino acid depletion relative to amino acid-replete state were plotted. *n* = 10713. *n* = 3 biological replicates for each condition. (c) Genes stabilized after amino acid depletion are related to metabolic processes and macromolecule degradation pathways. KEGG pathway enrichment analysis was performed on 3,433 genes that were highly stabilized after amino acid depletion (>4-fold change threshold, median 5.7-fold change).

We next asked if the mRNAs that show the highest stabilization are associated with specific cellular functions. To test this, we performed a KEGG pathway enrichment analysis, which showed categories relevant to cellular stress, including lysosome and endosome biology, which is relevant to autophagy and DNA repair-relevant pathways (such as base excision repair and homologous recombination) (**Fig. 2c**, **Supplementary Table 2**). Additionally, numerous categories related to amino acid and other metabolic pathways were enriched, potentially revealing pathways to overcome amino acid depletion. Overall, these results suggest that ISR-induced mRNA stabilization may contribute to homeostatic responses after stress.

### m^6^A levels correlate with transcript abundance increases seen in the ISR

We next wanted to understand the mechanism of mRNA stabilization during the ISR. Since a key feature of the ISR is reduced translation, we asked if translation-dependent mRNA degradation pathways might be impaired during the ISR. A major mechanism of translation-dependent mRNA degradation is m^6^A-mediated degradation^36^. We recently showed that m^6^A-mediated mRNA degradation is initiated when the translating ribosome encounters m^6^A in the coding region, impairing ribosome dynamics^36,38^. Thus, the reduced translation seen during the ISR could impair m^6^A-mRNA degradation.

As a first test, we asked if the mRNAs that show increased abundance during the ISR contain m^6^A and if the increased expression correlates with the amount of m^6^A in the transcript. To test this, we measured the change in mRNA abundance for each mRNA and grouped mRNAs based on their cumulative m^6^A levels. We quantified the cumulative m^6^A levels for each mRNA by summing the m^6^A stoichiometries at each m^6^A site in the transcript, based on the GLORI dataset^42^. This analysis showed that lowly methylated mRNAs showed reduced expression after amino acid depletion. However, the highly methylated mRNA showed increased expression after amino acid depletion. Importantly, the m^6^A levels directly correlated with increased mRNA expression (**Fig. 3a**, left). Overall, this suggests that activation of the ISR leads to suppression of the m^6^A-mediated mRNA decay pathway.

**Fig. 3:**
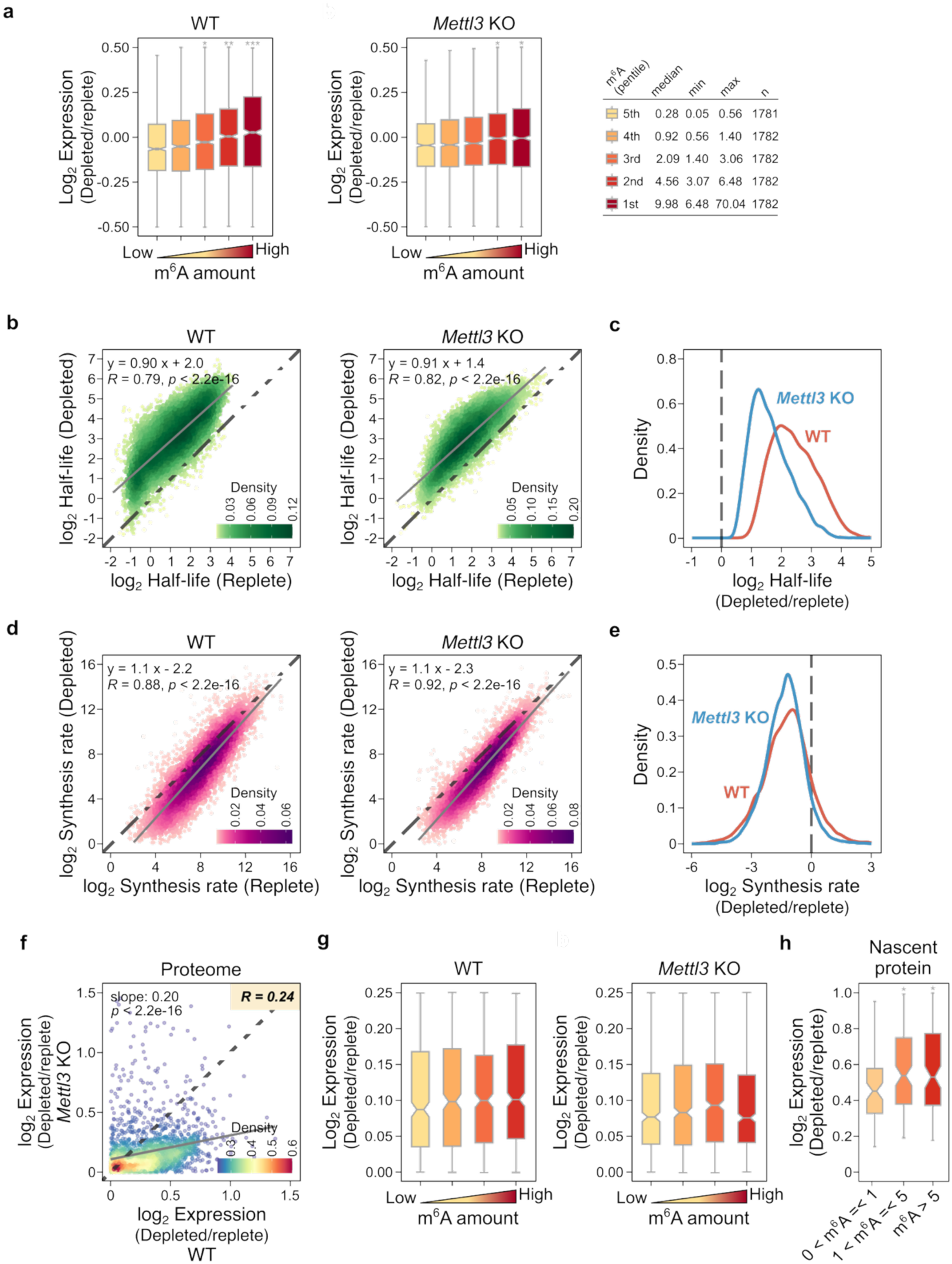
Global mRNA stabilization during the ISR depends on m^6^A. (a) m^6^A-mRNA expression increases after amino acid depletion. The mRNA expression levels were plotted in MEFs cultured in either amino acid-replete or -depleted media for 3 hours. mRNAs were grouped into pentiles based on their overall m^6^A levels determined by GLORI^42^. For each group of genes, we plotted the change in mRNA expression in the amino acid-depleted condition relative to the amino acid-replete condition. For lowly methylated genes, mRNA expression decreased after amino acid condition. However, highly methylated genes showed increased mRNA expression after amino acid condition. This effect was in proportion to the methylation levels of each mRNA. Two-sided Wilcoxon signed-rank test; *\*P < 2e-04, **P < 2e-08, ***P < 2e-12, ****P < 2e-16*. Each box shows the first quartile, median and third quartile, and whiskers represent 1.5× interquartile ranges. The table on the right indicates the minimum, median, and maximum cumulative amount of m^6^A per transcript and the number of transcripts for each pentile group. (b) Global increase in mRNA half-life during the ISR is in part mediated by m^6^A. (left) The same plot as in Fig. 2a, showing global increase in mRNA half-life after amino acid depletion in wildtype MEFs. (right) In *Mettl3* knockout MEFs, mRNAs are generally stabilized after amino acid depletion. However, the degree of stabilization was much smaller compared to their stabilization effects in wildtype MEFs. *n* = 10713 transcripts. *n* = 3 biological replicates for each condition. (c) Change in mRNA half-life after 3 h amino acid depletion relative to amino acid-replete condition in wildtype and *Mettl3* knockout MEFs were plotted. *n* = 10713. *n* = 3. (d) Global decrease in mRNA synthesis rates during the ISR is independent of m^6^A. (left) The same plot as in Fig. 1b, showing global reduction in mRNA synthesis after amino acid depletion in wildtype MEFs. (right) In *Mettl3* knockout MEFs, mRNA synthesis was similarly reduced. *n* = 10713 transcripts. *n* = 3 biological replicates for each condition. (e) Change in mRNA synthesis rates after 3 h amino acid depletion relative to the amino acid-replete condition in wildtype and *Mettl3* knockout MEFs was plotted. *n* = 10713. *n* = 3. (f) m^6^A mediates the proteomic changes in response to amino acid depletion. To test the role of m^6^A in the proteomic response to amino acid depletion, we measured protein expression changes in amino acid-depleted relative to amino acid-replete wild-type and *Mettl3* knockout MEFs. Cells were grown in amino acid-replete or amino acid-depleted media for 3 h, and protein expression was determined using label-free mass spectrometry. Amino acid depletion-mediated increase in protein expression was markedly reduced in *Mettl3* knockout MEFs relative to wild-type MEFs. This suggests that m^6^A plays a role in the proteomic response to amino acid depletion. (g) The increased expression of proteins during amino acid depletion directly correlates with the m^6^A amount in mRNAs that encode them. We quantified the protein expression induced during amino acid depletion by label-free mass spectrometry. We quantified increased expression of proteins based on the amount of m^6^A in the mRNAs that encode them. Two-sided Wilcoxon signed-rank test; *\*P < 2e-04, **P < 2e-08, ***P < 2e-12, ****P < 2e-16*. Each box shows the first quartile, median and third quartile, and whiskers represent 1.5× interquartile ranges. (h) m^6^A levels correlate with new protein synthesis during amino acid depletion. We analyzed a previously published nascent peptide-profiling dataset^45^ in which nascent proteins were measured using metabolic incorporation of azidohomoalanine in HeLa cells after 6 h of serum and amino acid depletion. m^6^A levels in mRNAs are correlated with the increased expression of corresponding proteins, consistent with the increased expression of m^6^A-mRNA and the corresponding protein synthesis rates after amino acid depletion. Two-sided Wilcoxon signed-rank test; *\*P < 2e-04, **P < 2e-08, ***P < 2e-12, ****P < 2e-16*. Each box shows the first quartile, median and third quartile, and whiskers represent 1.5× interquartile ranges.

A feature of m^6^A mRNAs is that they tend to be longer than other mRNAs and are derived from genes that contain long internal exons. These features act as triggers for co-transcriptional methylation. To rule out the possibility that these aspects of gene and mRNA structure might be regulated by the ISR, rather than m^6^A itself, we examined mRNA stabilization in *Mettl3* knockout MEFs. In *Mettl3* knockout MEFs the selective increase in mRNA abundance was markedly reduced (**Fig. 3a**). This suggests that m^6^A, rather than these co-occurring genomic features accounts for their stabilization after the ISR.

Together, these results suggests that ISR activation leads to suppression of the m^6^A-mediated mRNA decay pathway.

### ISR-induced transcriptome-wide changes in gene expression are markedly reduced in m^6^A-deficient cells

Although the ISR leads to stabilization of m^6^A mRNAs, we wanted to determine how much the increase in mRNA half-life of m^6^A-mRNAs contributes to the overall increase in mRNA half-life seen after amino acid depletion. To test this, we used *Mettl3* knockout MEFs and measured mRNA half-life measurement using TimeLapse-seq. In wild-type MEFs, amino acid depletion led to the expected global increase in mRNA half-lives (**Fig, 3b**, left). However, in *Mettl3* knockout MEFs, amino acid depletion led to much less prominent increase in mRNA half-life (**Fig. 3b** and **3c**). Although global mRNA stabilization was still detected, the degree of stabilization was much less than in wild-type cells. Notably, gene transcription induced by the ISR was maintained in *Mettl3* knockout MEFs (**Fig. 3d** and **3e**). Overall, these results indicate that impairment of m^6^A-mediated mRNA degradation mediates much of the increase in mRNA half-life after amino acid depletion, but there are also m^6^A-independent mechanisms of mRNA stabilization during the ISR.

### m^6^A-mRNA stabilization contributes to the overall proteomic landscape of the ISR

Although mRNA expression is found to change in the ISR, protein levels will mediate the final cellular response to stress. Although the ISR leads to a substantial reduction in translation, low levels of translation are maintained during the ISR^43,44^. We therefore hypothesized that mRNAs that are stabilized may contribute to new protein expression during the ISR.

We next asked if the proteomic changes seen in the ISR require m^6^A. To test this, we measured protein expression changes induced by amino acid depletion in both wild-type and *Mettl3* knockout MEFs using label-free proteomics. When we examined the proteins that increased in wild-type MEFs after amino acid depletion, we found that these proteins were generally much less increased in *Mettl3* knockout MEFs (**Fig. 3f**). These data suggest that the presence of m^6^A is needed for the increased protein expression.

Because m^6^A depletion can cause many indirect effects, we wanted to more directly examine protein expression from m^6^A-containing mRNAs. These mRNAs show increased expression in proportion to their m^6^A levels after amino acid depletion (see **Fig. 3a**, left). We therefore asked if their m^6^A levels correlates with the increase in protein levels from these mRNAs after amino acid depletion. For each protein, we quantified the cumulative m^6^A levels in the mRNAs that encode them by summing the m^6^A stoichiometries at each m^6^A site based on the GLORI dataset^42^. In wild-type cells, we found that the increase in protein levels correlated with the m^6^A levels on mRNAs that encode them (**Fig. 3g**, left). However, in *Mettl3* knockout MEFs, there was no clear correlation between increased protein expression and the m^6^A levels normally found on the mRNAs that encode them (**Fig. 3g**, right). These results suggest that the increased protein levels seen during the ISR is due, in part, to the stabilization of specific m^6^A-mRNAs, resulting in increased mRNA levels and a corresponding increase in protein levels.

We next wanted to determine if the same effect could be seen in datasets prepared by other labs. A recent study examined new protein synthesis induced by 6 h of amino acid and serum depletion in HeLa cells^45^. In these experiments, nascent proteins were labeled by incorporation of azidohomoalanine, a methionine analog, and the labeled proteins were enriched by pull-down for mass spectroscopy. Nascent proteins were measured in both amino acid-replete cells and in cells depleted of both serum and amino-acids. Using this dataset, we determined the fold-increase in protein synthesis for each of the detected proteins. Notably, lowly methylated mRNAs showed a relatively small increases in protein expression while more highly methylated mRNAs were associated with larger increases in protein expression (**Fig. 3h**). These data are consistent with our proteomic analysis in MEFs, and further support the idea that m^6^A-mRNA stabilization contributes to the altered protein expression seen in the ISR.

### m^6^A-mRNA stabilization is elicited by diverse inducers of the ISR

We next asked if other inducers of the ISR also increase the abundance of highly methylated mRNAs. To test this, we used published RNA-seq datasets obtained after treating cells with various inducers of the ISR. We first examined mRNA expression in U2OS cells after 2 h of thapsigargin treatment, which elicits endoplasmic reticulum stress. Here, we saw decreased expression for lowly methylated transcripts after thapsigargin treatment, while highly methylated transcript showed selective increase in their expression. Similar to amino acid depletion, the selective increase of highly methylated transcripts was in proportion to their methylation levels (**Fig. 4a**).

**Fig. 4:**
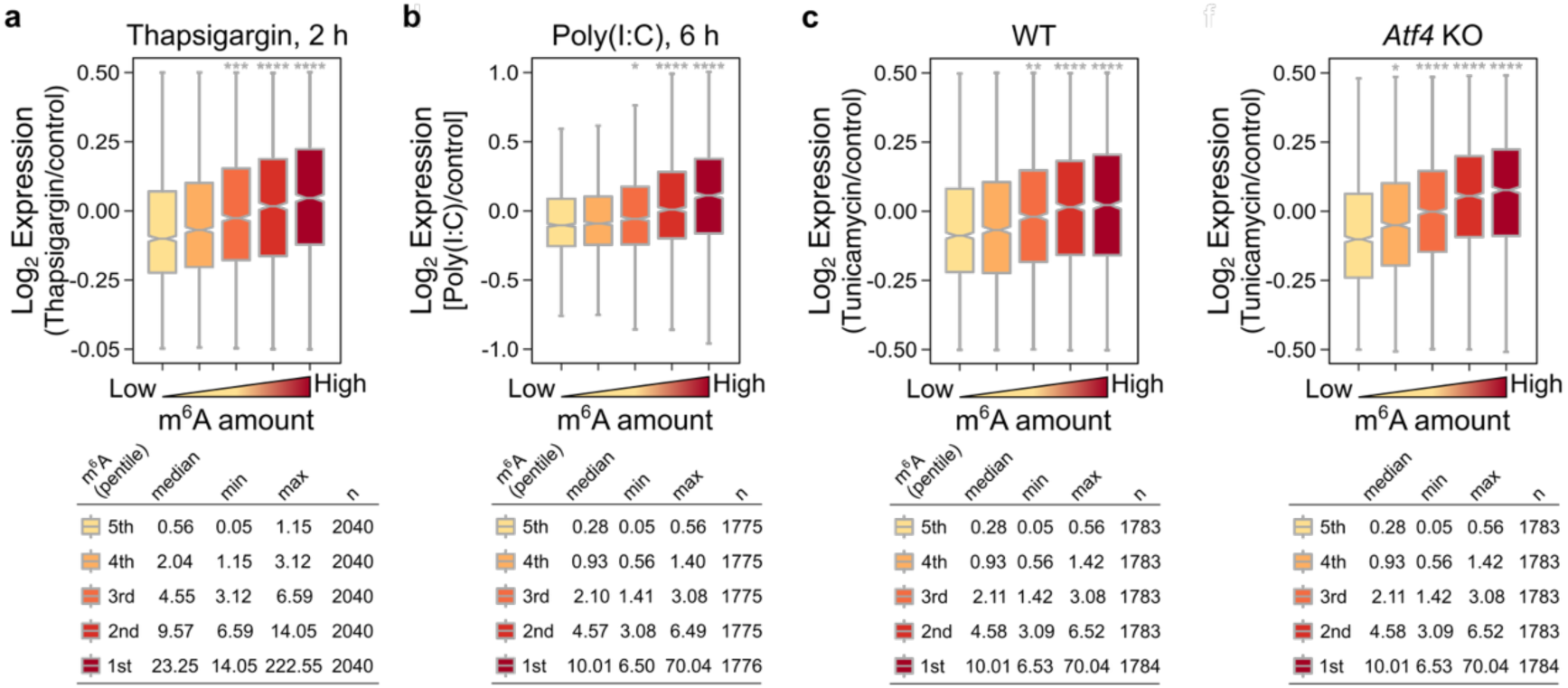
m^6^A-mRNA stabilization is elicited by diverse inducers of the ISR. (a) The ISR leads to the increase in m^6^A-mRNA expression. To test if other forms of cellular stress beside amino acid depletion maintain m^6^A-mRNA expression, we examined gene expression after other ISR inducers. The mRNA expression levels were plotted in U2OS cells after treatment with thapsigargin, an endoplasmic reticulum stress inducer of ISR, relative to vehicle treatment. mRNAs were grouped into pentiles based on m^6^A levels determined by GLORI^42^. Shown are box plots of the fold-change in gene expression after 2 h thapsigargin treatment relative to the control treatment. Highly methylated mRNAs showed increased expression after thapsigargin treatment relative to lowly methylated mRNAs. The data was obtained from GSE273600^80^. Two-sided Wilcoxon signed-rank test; *\*P < 2e-04, **P < 2e-08, ***P < 2e-12, ****P < 2e-16*. Each box shows the first quartile, median and third quartile, and whiskers represent 1.5× interquartile ranges. (b) The mRNA expression levels were plotted in MEF after transfecting poly(I:C), a double-stranded RNA that induces ISR, compared to vehicle treatment. mRNAs were grouped into pentiles based on m^6^A levels determined by GLORI^42^. Shown are box plots of the fold-change in gene expression 6 h after poly(I:C) transfection relative to the control mock transfection. Highly methylated mRNAs showed increased expression after poly(I:C) transfection relative to lowly methylated mRNAs. The data was obtained from GSE111938^81^. Two-sided Wilcoxon signed-rank test; *\*P < 2e-04, **P < 2e-08, ***P < 2e-12, ****P < 2e-16*. Each box shows the first quartile, median and third quartile, and whiskers represent 1.5× interquartile ranges. (c) To test if the increased expression of m^6^A-mRNA during ISR is independent of ATF4, we measured mRNA expression in the absence and presence of tunicamycin, an endoplasmic reticulum stress inducer of ISR, in the wild-type and *Atf4* knockout MEFs cells. mRNAs were grouped in pentiles based on their overall m^6^A levels determined by GLORI^42^. For each group of genes, we plotted the change in mRNA expression after tunicamycin treatment relative to vehicle treatment. For lowly methylated genes, mRNA expression decreased after tunicamycin treatment. However, highly methylated genes showed increased mRNA expression after tunicamycin treatment. This effect was in proportion to the amount of m^6^A in each mRNA and was similar in the wild-type and *Atf4* knockout MEFs. The data was obtained from GSE158605^17^. The plot for the wildtype (left) is a similar analysis to Figure S6D in Murakami *et al*.^36^ using the same dataset, and used here to compare with *Atf4* knockout cells. Two-sided Wilcoxon signed-rank test; *\*P < 2e-04, **P < 2e-08, ***P < 2e-12, ****P < 2e-16*. Each box shows the first quartile, median and third quartile, and whiskers represent 1.5× interquartile ranges.

We also examined datasets of MEFs treated with poly(I:C), an inducer of the ISR which mimics double-stranded RNA viral infection. Again, we saw a selective increase of highly methylated mRNA after poly(I:C) transfection in MEFs (**Fig. 4b**). Additionally, other inducers of the ISR, including hypoxia (24 h, HeLa cells), arsenite (2 h, U2OS), and heme depletion elicited by desferrioxamine treatment (6 h, MCF-7 cells), all showed similar trends but with smaller effect sizes (**Extended Data Fig. 1**). These findings support the idea that activation of the ISR through diverse stimuli, rather than specifically amino acid depletion, all lead to increases in m^6^A-mRNA transcript abundance.

Next, we wanted to rule out the possibility that the selective increase of highly methylated transcripts during the ISR is mediated by ATF4. To test this, we examined gene expression in control and *Atf4* knockout MEFs after endoplasmic reticulum stress induced by tunicamycin^17^. Tunicamycin induced a selective increase of highly methylated mRNAs, like the other ISR inducers (**Fig. 4c**, left). When we repeated the same analysis using *Atf4* knockout MEFs, the selective increase in expression of the highly methylated mRNAs was maintained (**Fig. 4c**, right). Thus, the increase in the expression of highly methylated m^6^A-mRNA is not mediated by ATF4.

Overall, these results suggest that m^6^A-mRNA stabilization is a general mechanism to control gene expression during the ISR, and this pathway is distinct from of ATF4-mediated gene expression.

### m^6^A inhibition leads to an ISR-like transcriptomic and proteomic response

m^6^A depletion is seen in diverse contexts, including various cancer models^46,47,48^ and neurons during neurodegeneration^49,50,51^. Additionally, m^6^A depletion by administration of METTL3 inhibitors is being tested in clinical trials as a therapeutic approach for specific cancers^52^. Since the ISR suppresses m^6^A-mediated degradation, we reasoned that m^6^A pathway inhibition in normal unstressed cells (amino acid-replete conditions) may induce gene and protein expression changes that normally occur with the ISR.

As described above, the ISR is associated with two different mechanisms to increase gene expression: (1) An increase in ATF4-dependent transcription; and (2) increased mRNA stability, due in part to suppression of the m^6^A-mediated degradation. The increase in expression of the ATF4 transcriptional network, in principle would not be induced by m^6^A depletion, since this effect is transcriptional, not post-transcriptional at the level of mRNA stability. However, when we examined ATF4 target genes in unstressed *Mettl3* knockout cells, we found that they exhibited higher expression levels of ATF4 target genes (**Fig. 5a**). This increase in ATF4 target genes does not appear to be due to ATF4-dependent transcription since we found no increase in eIF2alpha phosphorylation and no increase in ATF4 protein expression (**Fig. 5b and Extended Data Fig. 2a**).

**Fig. 5:**
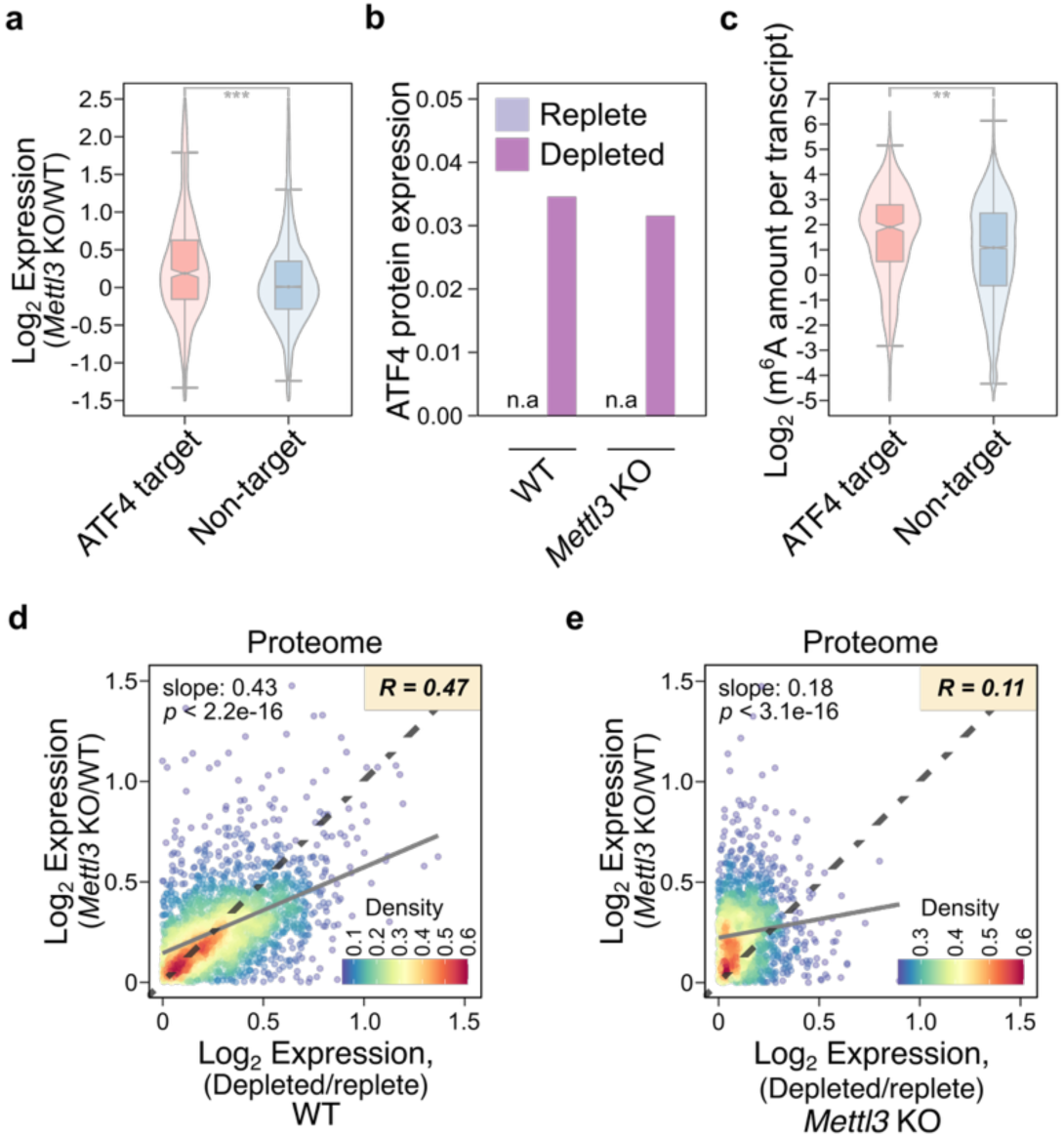
The transcriptome and proteome of m^6^A-depleted cells exhibit features of amino acid-depleted cells. (a) m^6^A depletion leads to the increased expression of ATF4 target genes. The increase in ATF4 target genes during the ISR is generally thought to be due to transcriptional activation. To test if the expression of these genes is affected by m^6^A-mRNA stabilization, we quantified their expression in *Mettl3* knockout MEFs relative to wildtype MEFs. Unexpectedly, we saw a preferential increase in expression of ATF4 target genes (n=564) compared to all other genes (non-target, n=10,144). Thus, ATF4 target genes are regulated through transcription activation and mRNA stabilization during the ISR. Two-sided Wilcoxon signed-rank test; *\*P < 2e-04, **P < 2e-08, ***P < 2e-12, ****P < 2e-16*. Each box shows the first quartile, median and third quartile, and whiskers represent 1.5× interquartile ranges. (b) m^6^A depletion does not affect ATF4 protein expression. The normalized ATF4 expression in label-free mass spectrometry data in wildtype and Mettl3 knockout MEFs under amino acid-replete and amino acid-depleted conditions. (c) ATF4 target genes are enriched with m^6^A. To ask whether the increased expression of ATF4 target genes in *Mettl3* knockout MEFs is a direct effect of m^6^A depletion, we quantified the amount of m^6^A in each transcript of ATF4 target genes (n=564) and all other genes (non-target, n=10,144). ATF4 target genes contain more m^6^A per transcript than all other genes expressed in MEFs. Two-sided Wilcoxon signed-rank test; *\*P < 2e-04, **P < 2e-08, ***P < 2e-12, ****P < 2e-16*. Each box shows the first quartile, median and third quartile, and whiskers represent 1.5× interquartile ranges. (d) m^6^A-depleted cells exhibit protein expression patterns that resemble amino acid-depleted cells. We measured protein expression changes in amino acid-replete *Mettl3* knockout MEFs relative to wild-type MEFs. To test the protein expression changes mediated by mRNA stabilization, we focused on the analysis on proteins that are induced after amino acid depletion. These m^6^A-mediated protein expression changes were correlated with the protein expression changes induced by amino acid depletion in wildtype MEFs. This suggests that suppression of m^6^A-mediated mRNA degradation contributes to the proteomic changes seen in response to amino acid-depletion. (e) To test if m^6^A determines the similarity in induced protein expression patterns between m^6^A-depleted cells and in amino acid-depleted state, we measured protein expression induced by amino acid depletion in *Mettl3* knockout MEFs. Amino acid depletion-mediated increase of protein expression was markedly reduced in *Mettl3* knockout cells. Protein expression increases seen in *Mettl3* knockout MEFs relative to the wild type MEFs showed poor correlation with amino acid-induced increases in protein expression changes in *Mettl3* knockout MEFs.

**Fig. 6:**
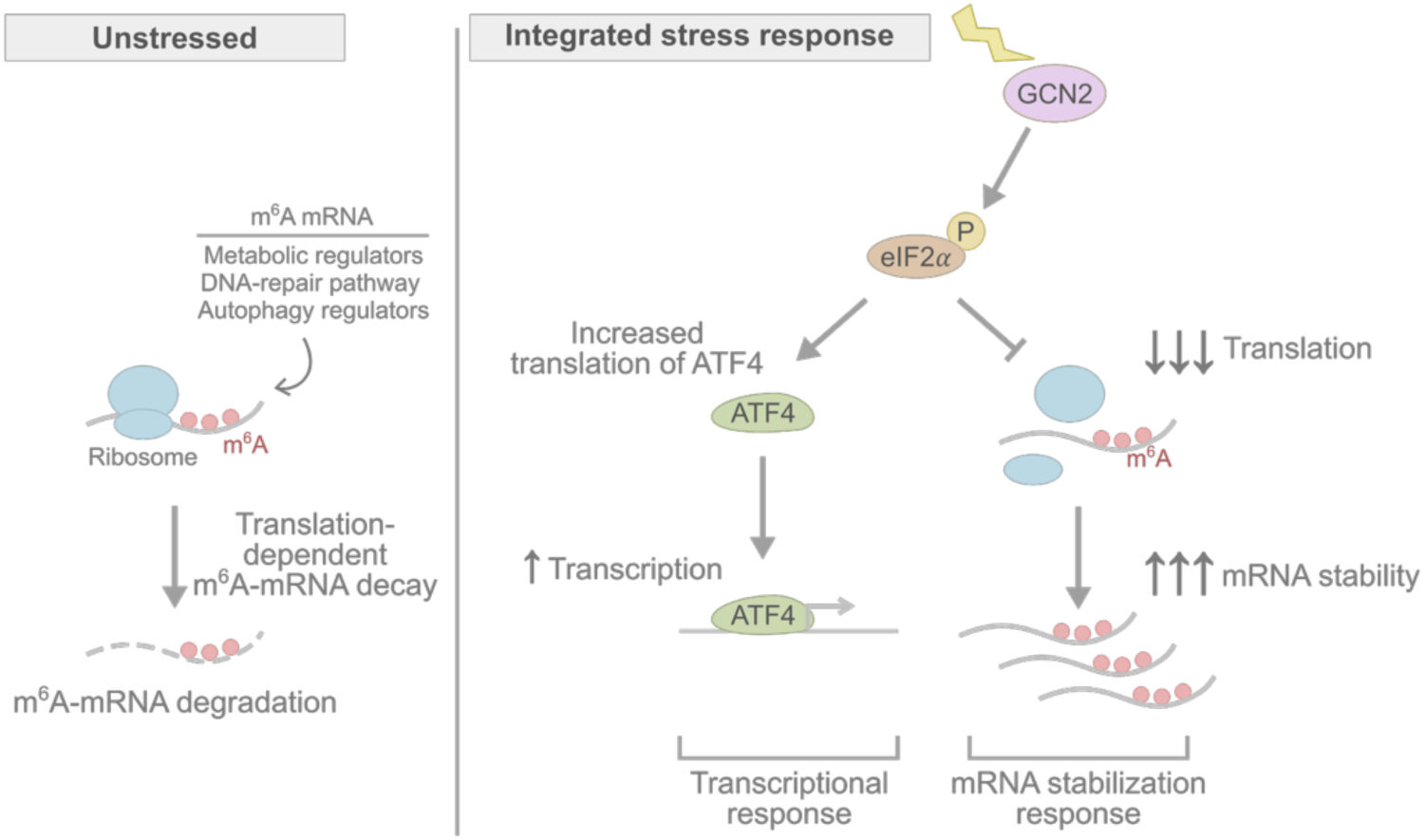
The ISR stabilizes m^6^A-mRNAs by repressing m^6^A-mediated mRNA decay. The ISR is a pathway that enables the cell to couple cellular stress to specific transcriptional responses. The cell expresses specific kinases that are activated upon different forms of stress, and then phosphorylate eIF2*α*, a critical regulator of translation. Upon phosphorylation, translation is impaired, which is generally thought to preserve amino acids. In the case of amino acid depletion, GCN2 is activated, which in turn phosphorylates eIF2*α*. eIF2*α* phosphorylation is a critical step in the ISR and leads to increased translation of ATF4, a master regulator of cellular stress responses. ATF4 activates the transcription of many important genes that contribute to the adaptive response to cell stress. Our data show that amino acid deprivation is associated with both new transcription and widespread mRNA stabilization. We show that the mRNA stabilization response is due to stabilization of m^6^A-mRNAs, many of which encode important regulators of stress response pathways such as autophagy. In unstressed conditions, where translation is active, highly methylated mRNAs are lowly expressed due to m^6^A-mediated mRNA degradation (left). However, upon eIF2*α* phosphorylation, translation is repressed, which leads to increased abundance of m^6^A-mRNAs. This effect is due to an essential role for translation in m^6^A-mediated mRNA degradation.Although translation is reduced in amino acid-depleted cells, the abundance of highly methylated mRNA is increased sufficiently to lead to an overall increase in protein expression from these transcripts. The increased protein expression from these highly methylated transcripts coordinates the adaptive cellular responses such as autophagy induction. Overall, the ISR, and specifically eIF2*α* phosphorylation, coordinates both new transcription and m^6^A-mRNA stabilization to mediate the adaptive response to stress.

To explain the upregulation of ATF4 target genes, we considered the possibility that they were stabilized by *Mettl3* knockout, rather than transcriptionally induced. To test this, we examined whether ATF4 target genes are more methylated than other genes in the cell. Here we found a clear increased level of m^6^A in ATF4 target genes (**Fig. 5c**). Additionally, we found that ATF4 target genes showed a larger increase in mRNA half-life after METTL3 inhibition than other genes in the cell (**Extended Data Fig. 2b**).

We next asked if m^6^A depletion induces a proteomic response that resembles the changes in the proteome elicited by amino acid depletion. To test this, we used label-free proteomics to measure protein expression changes induced by m^6^A depletion in *Mettl3* knockout MEFs, and we then compared the result with protein expression changes seen during the ISR induced by amino acid depletion in wild-type MEFs. For proteins that exhibit increased expression after amino acid depletion, we saw a general correlation between their expression and protein expression increases seen upon *Mettl3* knockout (**Fig. 5d** and **5e**). Thus, m^6^A depletion leads to a proteomic signature that has features of the ISR proteome elicited by amino acid-depleted states.

Overall, these data suggest that ATF4 target genes have higher m^6^A levels, thus likely contributing to their low expression in basal conditions, but their expression is induced by both transcriptional upregulation and reduced m^6^A-mediated degradation in stress. *METTL3* inhibition is therefore sufficient to induce many ATF4 target genes without a typical stressor.

## DISCUSSION

It is well known that the ISR leads to transcriptomic alterations^53,54^, and this effect is thought to be mediated by transcriptional upregulation by ATF4^2,15^. Although ATF4-mediated transcription clearly mediates key aspects of the cellular responses to stress^2^, ATF4-mediated transcription only accounts for ∼7.5% to ∼36% of the overall gene expression changes seen during cell stress^16,17^. Our TimeLapse-seq analysis shows that the transcriptomic response to stress involves both new transcription and mRNA stabilization, with mRNA stabilization affecting a much larger number of transcripts and accounting for the majority of the increased transcript levels seen during stress. This mRNA stabilization is mediated by suppression of the m^6^A mRNA degradation pathway. In basal conditions, m^6^A mRNAs are actively degraded in a translation-dependent manner. However, the ISR leads to decreased translation, which impairs m^6^A-mediated degradation, resulting in increased levels of m^6^A-containing mRNAs. These stabilized mRNAs comprise a cohort of mRNAs linked to major adaptive responses to stress.

Overall, our study identifies m^6^A suppression as a major mechanism by which the ISR achieves transcriptomic changes that define the adaptive response to cell stress.

Although the ISR leads to increased expression of m^6^A-mRNAs, the impact of the increased expression might be limited since mRNA translation is generally reduced during the ISR. However, cell stresses reduce rather than completely block translation^43,44^, as we confirmed in our puromycin nascent protein labeling experiment (**Extended Data Fig. 2a**). Thus, even though translation rates are reduced, new proteins can still be synthesized, especially if the abundance of a transcript increases. Indeed, our proteomic analysis shows that the increased expression of proteins during the ISR correlates with level of m^6^A in the corresponding transcripts. m^6^A-mRNA stabilization may also be important after stress resolution, when translation resumes. In this case, protein synthesis will be biased towards the m^6^A-mRNAs that have accumulated as a result of m^6^A pathway suppression.

A previous study suggested that m^6^A in the *ATF4* mRNA may enhance its translation rate during stress^55^. However subsequent studies showed that the m^6^A mapped to *ATF4* was instead a related nucleotide (*N*^6^, 2’-*O*-dimethyladenosine)^56^. Indeed, we found no clear change in ATF4 protein levels in *Mettl3* knockout MEFs.

Numerous studies have shown that m^6^A levels are reduced in a variety of cancers^46,47^. m^6^A depletion increases their proliferation, metastasis, and therapeutic resistance^57,58,59,60,61,62,63^. Since m^6^A-containing mRNAs are stabilized in both the ISR and after m^6^A depletion seen in cancers, both contexts have similar induction of genes related to stress adaptation. Furthermore, we also found that many ATF4 target transcripts were enriched in m^6^A, and these mRNAs were stabilized by m^6^A depletion even in the absence of a cellular stress. Thus, m^6^A deficiency may be associated with a transcriptomic state that resembles features of the ISR. It is possible that the decreased m^6^A levels in certain cancers endows these cells with stress-related gene expression patterns that may contribute to their enhanced growth and proliferation.

## METHODS

### Cell culture

MEFs were grown at 37°C in 5% CO_2_ using DMEM (ThermoFisher, 11995065) supplemented with 10% FBS and 10 U/mL penicillin-streptomycin (ThermoFisher, 15140122). Cells were passaged every 3-4 days using with TrypLE Express (ThermoFisuer, 12604039) according to the manufacturer’s instructions.

To deplete amino acids, cells were washed with PBS (ThermoFisher, 10010049) three times, then the PBS from the last wash was replaced with amino acid-replete or amino acid-depleted media for 3 h. Amino acid-replete media was prepared using DMEM (US Biological, D9800) supplemented with 3.5 g/L glucose, 3.7g/L sodium bicarbonate, and 10% dialyzed FBS (ThermoFisher A3382001). The amino acid composition of this DMEM is: L-Arginine•HCl (0.084 g/L, 398.7 µM), L-Cystine•2HCl (0.0626 g/L, 199.9 µM), L-Glutamine (0.584 g/L, 4.0 mM), Glycine (0.03 g/L, 399.6 µM), L-Histidine•HCl•H_2_O (0.042 g/L, 200.4 µM), L-Isoleucine (0.105 g/L, 800.5 µM), L-Leucine (0.105 g/L, 800.5 µM), L-Lysine•HCl (0.146 g/L, 998.7 µM), L-Methionine (0.03 g/L, 201.1 µM), L-Phenylalanine (0.066 g/L, 399.5 µM), L-Serine (0.042 g/L, 399.7 µM), L-Threonine (0.095 g/L, 797.5 µM), L-Tryptophan (0.016 g/L, 78.3 µM), L-Tyrosine•2Na•2H_2_O (0.10379 g/L, 397.3 µM), L-Valine (0.094 g/L, 802.3 µM). Amino acid-depleted media was prepared using amino acid-free DMEM (US Biological, D9800-13) supplemented with 3.5 g/L glucose, 3.7 g/L sodium bicarbonate, 1 mM sodium pyruvate, and 10% dialyzed FBS (ThermoFisher, A3382001).

### RNA-seq analysis

Adapters in RNA-seq reads were first trimmed using TrimGalore-0.6.9 with the default parameters. For the paired-end libraries, --paired option was added. After adapter trimming, read quality was assessed using FastQC v0.11.9 and Multiqc, v1.14 and reads were quantified using Salmon v1.5.2 using the default parameters. The read quantification was summarized at gene-level using tixmport package^64,65^. To analyze differential expression, the gene-level quantification result was performed using DEseq2 v1.34.0 with the default parameters^66^. Further computation and plotting were performed using a custom script on R v4.1.2^67^.

### TimeLapse-seq library preparation and sequencing

TimeLapse-seq was performed as previously described^41^. Briefly, cells were grown to 80% confluency. Cells were washed with PBS for three times and then incubated in amino acid replete or amino acid-depleted media for 3 h with 500 µM 4-thiouridine (4sU, SigmaAldrich, T4509). After 3 h, total RNA was collected using Qiagen RNeasy Mini kit RLT buffer (Qiagen, 74104) supplemented with 1% *β*-mercaptoethanol (BME). Cells were lysed by passing through a 22-gauge needle, and RNA was isolated following the manufacturer’s instruction with RPE buffer supplemented with 1% BME. 10 µg total RNA was treated with 2,2,2-trifluoroethylamine (TFEE) and sodium periodate (NaIO_4_) at 45 °C for 1 h to convert 4sU to *N*^4^-trifluoroethylcytosine. RNA was then purified twice using RNAClean beads (Beckman Coulter, A63987). 1 µg of treated RNA was used for RNA-seq library preparation using NEBNext^®^ rRNA Depletion Kit (NEB, E6310X) and NEBNext^®^ Ultra^™^ II Directional RNA Library Prep Kit (NEB, E7760L). Library quality was assessed on Agilent TapeStation4200 using High Sensitivity DNA ScreenTape (Agilent, 5067-5584). Library was quantified on Qubit4 Fluorometer using Qubit 1X dsDNA HS Assay Kit (ThermoFisher, Q33231). Libraries were sequenced with pair-end 2×100 cycles on Illumina NovaSeq 6000.

### TimeLapse-seq library data analysis

Library quality was first analyzed using MultiQC 1.14^68^. Adaptors were trimmed using TrimGalore-0.6.9^69^ with --stringency 3 and library quality was again assessed using MultiQC 1.14^68^ The data was then analyzed using the slamdunk all function of the analytical suite SLAM-DUNK 0.4.3 with the default parameters for mapping, quality filtering, and single nucleotide polymorphism identification.^70^ The reference genome hg38 and mm10 was used for mapping and the subsequent analysis. PCR duplicates in the quality filtered reads were collapsed using alleyoop collapse function of SLAM-DUNK 0.4.3. The 4sU conversion was further assessed using alleyoop rates, alleyoop tccontext, and alleyoop snpeval functions with the default parameters. alleyoop utrrates was used with a bed file of all exons of each mRNA to analyzed 4sU conversion rates per mRNA. The quality filtered and PCR duplicate-collapsed mapped reads were further analyzed using GRAND-SLAM 2.0.7b^71,72^ with the default parameters to quantify read counts and perform statistical analysis to infer the ratio of transcripts for each gene that originate from pre-4sU labeling and post labeling. The results from GRAND-SLAM 2.0.7b was used to calculate mRNA synthesis rate and half-life using grandR 0.2.2^73^. Further computation and plotting were performed using a custom script on R v4.1.2^67^.

### m^6^A quantification

We used the processed data deposited under each accession number for previously published GLORI datasets (**Supplementary Table 3**). The mapped m^6^A sites in the chromosome coordinates were first converted to the transcriptome coordinates using MetaPlotR^74^. The longest transcript for each gene was selected for the further analysis. For the GLORI dataset, the stoichiometry for each m^6^A site was provided appended to each m^6^A site at each transcriptome coordinate. Cumulative m^6^A amounts for 5’UTR, CDS, 3’UTR, and the entire transcript for each gene was calculated based on the output from MetaPlotR^74^ using a custom script on R v4.1.2^67^.

### Defining ATF4 target genes

We used previously published datasets to define ATF4 target genes. To define the set of ATF4 target genes used in Fig. 1d, we used the previously published ChIP-seq dataset (GSE35681)^75^. In this dataset, genes with ATF4 binding sites within 3 kb from their promoters are defined as ATF4 target genes^75^. In **Fig. 5a**, **5c**, and **Extended Fig 2b**, ATF4 target genes are defined as genes whose expression increased during the ISR in wildtype MEFs, but not in *Atf4* knockout MEFs^17^(GSE158605)^17^.

### Proteomics

Cells were washed twice with ice-cold PBS on ice. Cells were scraped in 1 mL PBS and pelleted by centrifugation at 1000 x g for 5 min. The cells were lysed in 4% SDS and the protein was acetone precipitated and re-suspended in 0.1% RapiGest (Waters) containing 25 mM ammonium bicarbonate. The samples were then reduced with DTT, alkylated with iodoacetamide, and digested overnight with trypsin at 37 °C. The digests were desalted by C18 Stage-tip columns.

The digests were analyzed using a Thermo Fisher Scientific EASY-nLC 1200 coupled on-line to a Fusion Lumos mass spectrometer (Thermo Fisher Scientific). Buffer A (0.1% FA in water) and buffer B (0.1% FA in 80 % ACN) were used as mobile phases for gradient separation. A 75 µm x 15 cm chromatography column (ReproSil-Pur C18-AQ, 3 µm, Dr. Maisch GmbH, German) was packed in-house for peptide separation. Peptides were separated using a gradient of 5-40% buffer B over 30 min, 40-100% B over 10 min at a flow rate of 400 nL/min. The Fusion Lumos mass spectrometer was operated in a data independent acquisition (DIA) mode. MS1 scans were collected in the Orbitrap mass analyzer from 350-1400 m/z at 120K resolutions with the normalized automatic gain control (AGC) target set to 750 and a maximum ion accumulation time of 50 ms. The instrument was set to select precursors in 45 x 14 m/z wide windows with 1 m/z overlap from 350-975 m/z for HCD fragmentation. The MS/MS scans were collected in the orbitrap at 15K resolution. The normalized AGC setting was 6000 and the maximum ion accumulation time was 22 ms.

Data were analyzed, searched, filtered, and assembled into protein quantification values using DIA-NN v1.8^76^ with the following settings: protease: Trypsin/P, missed cleavages: 1, peptide length range 7-30, FDR: 1%, match between runs: enable, quantification strategy: “Robust LC (high precision)”, cross-run normalization: RT-dependent. The spectral library was generated in DIA-NN using in silico deep-learning and RT prediction based on the mouse Uniprot database entries (8/7/2021).

### KEGG pathway analysis

A list of genes was generated by filtering for the > 4-fold increase in mRNA half-life after 3 h amino acid depletion in MEFs measured using TimeLapse-seq (**Supplementary Table 2**). Using this gene list as the input, the KEGG pathway enrichment analysis was performed using enrichKEGG function using clusterProfiler 4.8.1^77,78^ on R version 4.3.0.

## DATA AVAILABILITY

Sequencing datasets generated in this study are deposited to NCBI Gene Expression Omnibus GSE249256. Accession numbers for previously published sequencing datasets used in this study are provided in **Supplementary Table 3**. The reference genome GRCh38 and GRCm39 was downloaded from https://useast.ensembl.org/index.html. The mass spectrometry proteomics data is available at https://massive.ucsd.edu/ProteoSAFe/static/massive.jsp using the login credential MSV000093588_reviewer. Additional information for the analyses of the data in this study is available upon request to the corresponding author.

## CODE AVAILABILITY

Original codes used in this study are available upon request to the corresponding author.

## ACKNOWLEDGMENTS

We thank J. G. Dumelie, V. Despic, M. Chen, M. Oleynikov, and the rest of members in Jaffrey laboratory for helpful comments and discussions. We also thank Genomics Resources Core Facility and Proteomics & Metabolomics Core Facility at Weill Cornell Medicine for providing illumine sequencing service and mass spectrometry service, respectively Supported by NIH grants S10 OD030335, RM1HG011563, and R35 NS111631 (S.R.J.), and Moderna Research Fellowship and Department of Defense fellowship BC180715 (S.M).

## AUTHOR INFORMATION

### Contributions

S.M. and S.R.J. conceived the project and designed the experiments. S.M. performed all experiments and analysis. S.M. and S.R.J. wrote the manuscript.

## SUPPLEMENTARY INFORMATION

### Supplementary Information

Supplementary Table 1 Differential expression in MEF after amino acid depletion (related to Fig. 1).

Supplementary Table 2 List of genes for KEGG pathway enrichment analysis.

Supplementary Table 3 List of previously published datasets used in this study.

---

**Extended Data Fig. 1:**
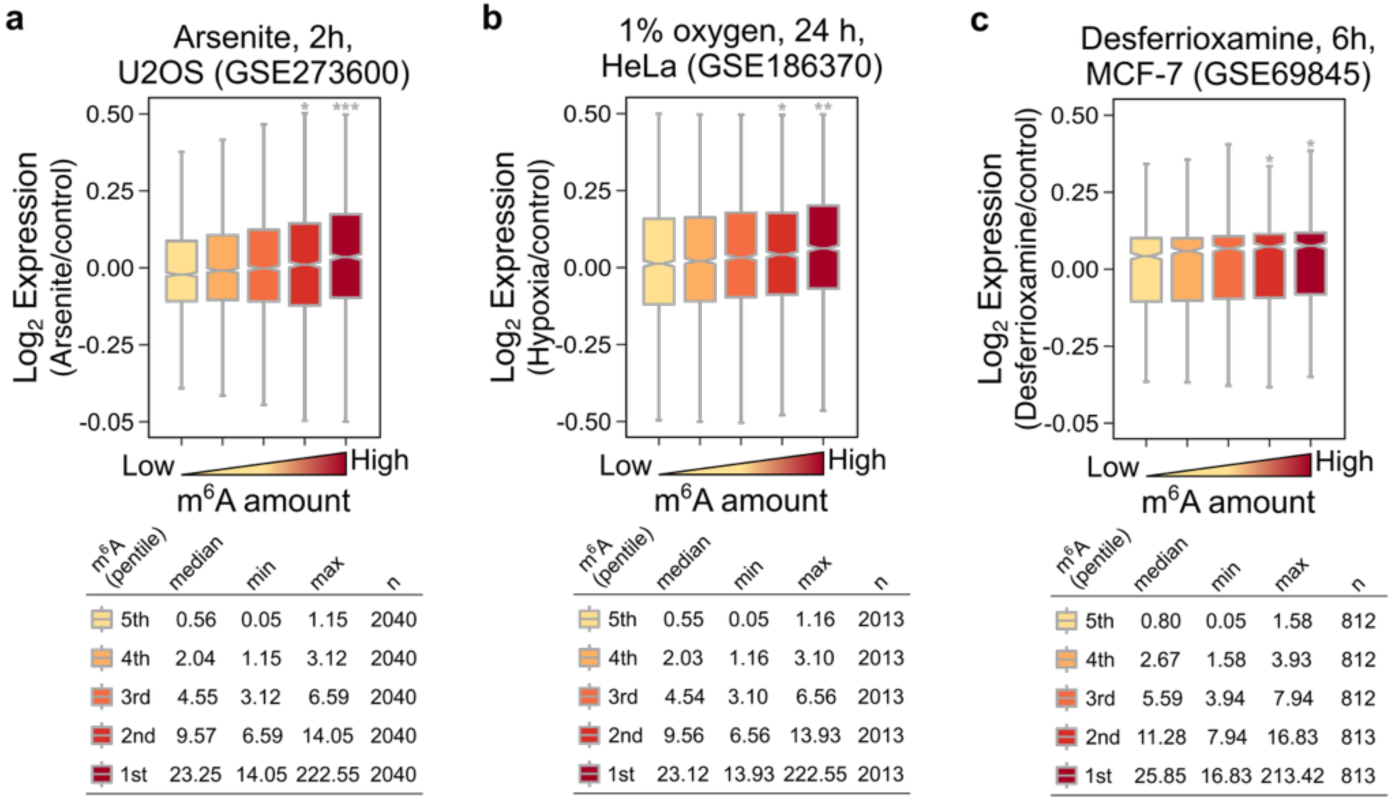
ISR selectively increases the abundance of highly methylated mRNA. (a) The mRNA expression levels were plotted in U2OS cells after treatment with arsenite, an oxidative stress inducer of ISR. mRNAs were grouped into pentiles based on m^6^A levels determined by GLORI^42^. Shown are box plots of the fold-change in gene expression after 2 h arsenite treatment relative to the control treatment. Highly methylated mRNAs showed increased expression levels after arsenite treatment relative to lowly methylated mRNAs. The data was obtained from GSE273600. Two-sided Wilcoxon signed-rank test; *\*P < 2e-04, **P < 2e-08, ***P < 2e-12, ****P < 2e-16*. Each box shows the first quartile, median and third quartile, and whiskers represent 1.5× interquartile ranges. (b) The mRNA expression levels were plotted in HeLa cells after exposure to hypoxia, an oxidative stress inducer of ISR. mRNAs were grouped into pentiles based on m^6^A levels determined by GLORI^42^. Shown are box plots of the fold-change in gene expression after culturing HeLa cells in hypoxic conditions for 2 h relative to the control normoxic condition. Highly methylated mRNAs showed increased expression levels in the hypoxic condition relative to lowly methylated mRNAs. The data was obtained from GSE186370. Two-sided Wilcoxon signed-rank test; *\*P < 2e-04, **P < 2e-08, ***P < 2e-12, ****P < 2e-16*. Each box shows the first quartile, median and third quartile, and whiskers represent 1.5× interquartile ranges. (c) The mRNA expression levels measured by microarray were plotted in MCF-7 cells after exposure to desferrioxamine that chelates iron leading to impaired heme synthesis. mRNAs were grouped into pentiles based on m^6^A levels determined by GLORI^42^. Shown are box plots of the fold-change in gene expression after culturing MCF-7 cells in desferrioxamine for 6 h relative to the control condition. Highly methylated mRNAs showed increased expression levels during heme depletion relative to lowly methylated mRNAs. The data was obtained from GSE186370. Two-sided Wilcoxon signed-rank test; *\*P < 2e-04, **P < 2e-08, ***P < 2e-12, ****P < 2e-16*. Each box shows the first quartile, median and third quartile, and whiskers represent 1.5× interquartile ranges.

**Extended Data Fig. 2:**
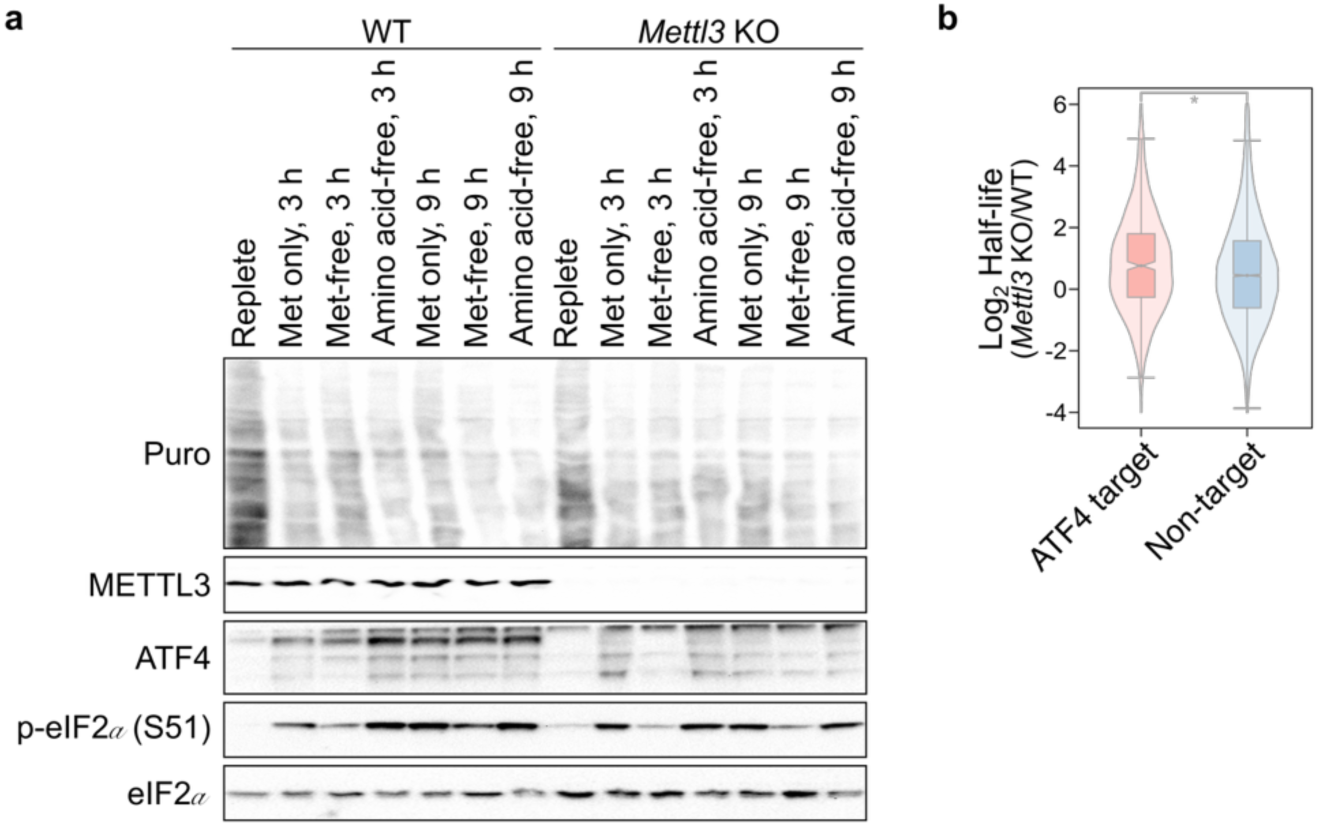
m^6^A depletion does not lead to eIF2*α* phosphorylation or ATF4 expression. (a) m^6^A depletion does not induce the phosphorylation of eIF2*α* or ATF4. Western blot showing the puromycin incorporation rates and the expression of METTL3, ATF4, and the phosphorylation of eIF2*α* under the indicated conditions in wildtype and *Mettl3* knockout MEFs. (b) m^6^A depletion leads to stabilization of ATF4 target genes. We asked whether the increased expression of ATF4 target genes after m^6^A depletion seen in Fig. 5a is due to mRNA stabilization. We examined mRNA half-lives of ATF4 target genes (n=564) and all other genes (non-target, n=10,144) in *Mettl3* knockout cells compared to wildtype MEFs. ATF4 target genes are more stabilized after m^6^A depletion compared to all other genes. Two-sided Wilcoxon signed-rank test; *\*P < 2e-04*. Each box shows the first quartile, median and third quartile, and whiskers represent 1.5× interquartile ranges.

